# Single-cell transcriptomes identify patient-tailored therapies for selective co-inhibition of cancer clones

**DOI:** 10.1101/2023.06.26.546571

**Authors:** Aleksandr Ianevski, Kristen Nader, Daria Bulanova, Anil K Giri, Tanja Ruokoranta, Heikki Kuusanmäki, Nemo Ikonen, Philipp Sergeev, Markus Vähä-Koskela, Anna Vähärautio, Mika Kontro, Kimmo Porkka, Caroline A. Heckman, Krister Wennerberg, Tero Aittokallio

## Abstract

Intratumoral cellular heterogeneity necessitates multi-targeting therapies for improved clinical benefits in patients with advanced malignancies. However, systematic identification of patient-specific treatments that selectively co-inhibit cancerous cell populations poses a combinatorial challenge, since the number of possible drug-dose combinations vastly exceeds what could be tested in scarce patient cells. Here, we developed scTherapy, a machine learning model that leverages single-cell transcriptomic profiles to prioritize multi-targeting treatment options for individual patients with hematological cancers or solid tumors.

## Introduction

High intratumoral heterogeneity and evolution of cancer cell populations are major drivers of therapy resistance both in hematological malignancies and solid tumors^1–5^. In acute myeloid leukemia (AML), a number of single-cell genomic analyses have mapped the clonal evolutionary processes of disease progression and therapy resistance at cell subpopulation level, as well as deciphered cellular hierarchy and reprogramming among leukemic cell subpopulations involved in the chemoresistance, relapse and clinical outcomes^6–9^. Similarly in solid tumors, clonal analysis and longitudinal sampling of patients with high-grade serous ovarian carcinoma (HGSC) have revealed evolutionary trajectories, with distinct genomic and morphological features that associate with treatment responses^10^. Despite this wealth of information, we lack approaches to target chemoresistant subpopulations to enhance second-line treatment efficacy in relapsed patients, or to avoid resistance to first-line therapies by co-inhibiting multiple leukemic cell subpopulations with sufficiently high potency and precision. There is a medical need for systematic approaches to identify more effective combinatorial therapies, using either multi-targeting inhibitors or their combinations, which selectively co-inhibit multiple signaling pathways that drive the disease-or resistance in heterogeneous patient and cell populations. Several computational approaches for *in silico* prediction of drug combination effects have been developed, yet we lack approaches that consider both the patient and disease heterogeneity when predicting drug sensitivity differences among cell populations, toward designing cancer-selective and patient-specific therapeutic options using feasible measurements in scarce patient cells.

To address these limitations, we implemented a machine learning model, scTherapy, which identifies cancer-selective and low-toxic multi-targeting options for each individual cancer patient based on scRNA-seq data alone. The selective predictions come from transcriptomic differences between genetically distinct cancer cell populations (or clones) in individual patient samples, when compared to non-cancerous cells from the same patient sample (**Fig. 1, Online Methods**). To enable fast translational applications, we pre-trained a gradient boosting model (LightGBM) that predicts drug response differences across cell populations by leveraging a massive reference database of large-scale phenotypic profiles (both transcriptomics and viability readouts) measured in cancer cell lines in response to single-drug perturbations. When applied to patient samples, the model generates a ranked list of most effective multi-targeting options (either targeted-agents, chemotherapies, or their combinations) that selectively co-inhibit key cancer clones in a given patient sample. To guide translational applications, we further remove low-confidence predictions and likely non-tolerated doses among the dose-specific drug response predictions, hence ensuring that only the most relevant predictions will be suggested for treatment optimization. The ScTherapy predictions makes functional *ex vivo* drug testing in patient-derived cells more feasible in a translational setting by prioritizing most potent multi-targeting options for further experimental validation in scarce patient cells. In doing so, we also extend the combinatorial space of single-cell drug response assays, which is currently constrained both by the patient cell availability and excessive time and cost of the assays for translational use.

**Fig. 1.**
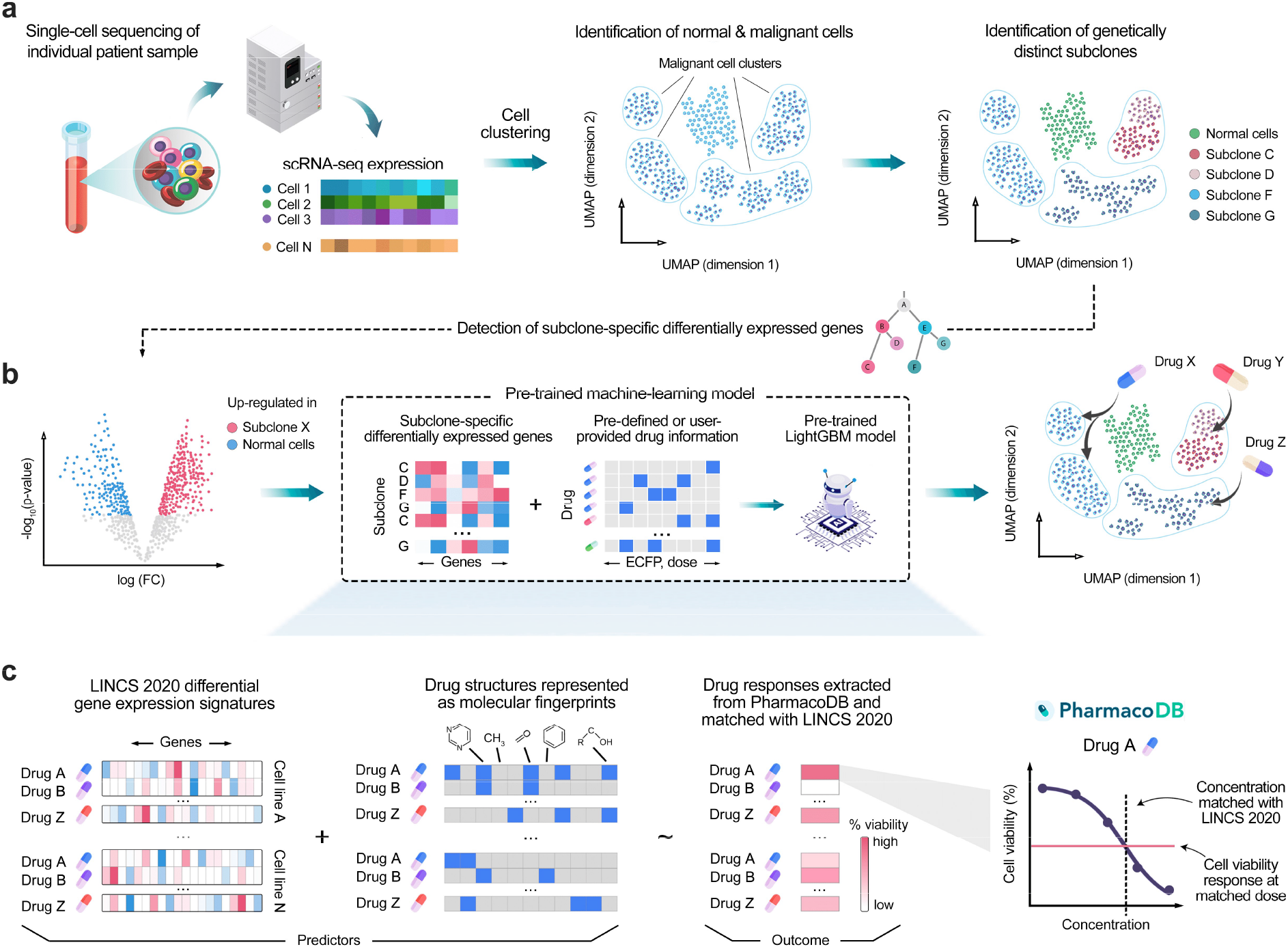
Schematic illustration of the experimental-computational prediction approach. Identification of clone-specific and cancer-selective compounds is performed in two steps: (**a**) Raw sequencing data from selected tissue are processed and aligned to generate a scRNA-seq expression count matrix. Unsupervised clustering separates malignant and normal cell clusters using an ensemble prediction approach with three analytical tools: ScType, CopyKAT and SCEVAN (**Suppl. Fig. 1**). InferCNV infers large-scale copy number variations and identifies genetically distinct subclones among the malignant cells. (**b**) Subsequently, subclone-specific differentially-expressed genes are identified through differential expression analysis. The identified genes, along with drug information such as molecular fingerprints and drug doses, serve as inputs for the pre-trained LightGBM model. Based on the patient-specific inputs, the pre-trained model predicts the most potent compounds and their effective doses for each subclone. (**c**) To train the LightGBM model, a comprehensive dataset was compiled that integrates transcriptional changes from small-molecule perturbation experiments (LINCS 2020 dataset), with chemical structures represented as ECFP fingerprints and drug-dose response data collected from various studies (PharmacoDB resource). Concentrations of the LINCS 2020 dataset were matched with dose-response curves from the PharmacoDB, and interpolated cell viability was used as the outcome variable for LightGBM model.

## Results

We developed the scTherapy model and tested its translational potential first by analyzing single-cell transcriptomic profiles of 12 bone marrow samples from diagnostic and refractory or relapsed AML patients (**Suppl. Table 1**), followed by careful experimental validation of the model predictions in the primary cells of the same patient samples.

To design multi-clone targeting and cancer-selective therapeutic options for each patient, we leveraged 394,303 genome-wide transcriptomic profiles post-treatment with 19,646 single-agent responses, measured in multiple doses in 167 cell lines, available from the LINCS 2020 project^11^. We next matched these transcriptomic response profiles with drug-induced cell viability responses available from PharmacoDB^12^, measured in multiple doses in the same 167 cell lines to pre-train a LightGBM that predicts drug response differences across cell populations (**Fig. 1**). The model predicts drug response using fold changes of differentially expressed genes (DEGs) after drug treatment at a particular dose, hence leading to concentration-specific cell inhibition predictions. In the patient applications, we used the pre-trained model to predict multi-targeting options that can selectively co-inhibit multiple cancer subclones, identified from patient-specific scRNA data, and using fold changes of DEGs between normal cells and cancer cell populations as input. In the final step, we combined the top-predicted effective and selective drugs for each clone as a targeted combinatorial therapy for the patient sample. This is the first translational approach for systematic tailoring of personalized multi-targeting options that takes into account both the intratumoral cellular heterogeneity and dose-specific therapeutic and toxic effects of anticancer compounds.

### Experimental validation of the model predictions in AML patient samples *ex vivo*

We carried out patient-specific treatment predictions using scRNA-seq profiles from bone marrow aspirates of 12 AML patients (**Suppl. Table 1**). The single-cell transcriptomes revealed highly heterogeneous cell type compositions across the patients, both for leukemic and normal cell types (**Fig. 2a**), necessitating personalized treatment predictions. Through processing the scRNA-seq data from each patient separately, and then feeding it into the ScTherapy model, we generated personalized predictions of drug treatments that are likely to be the most or least effective for each patient case (see **Online Methods** for more details).

**Fig. 2.**
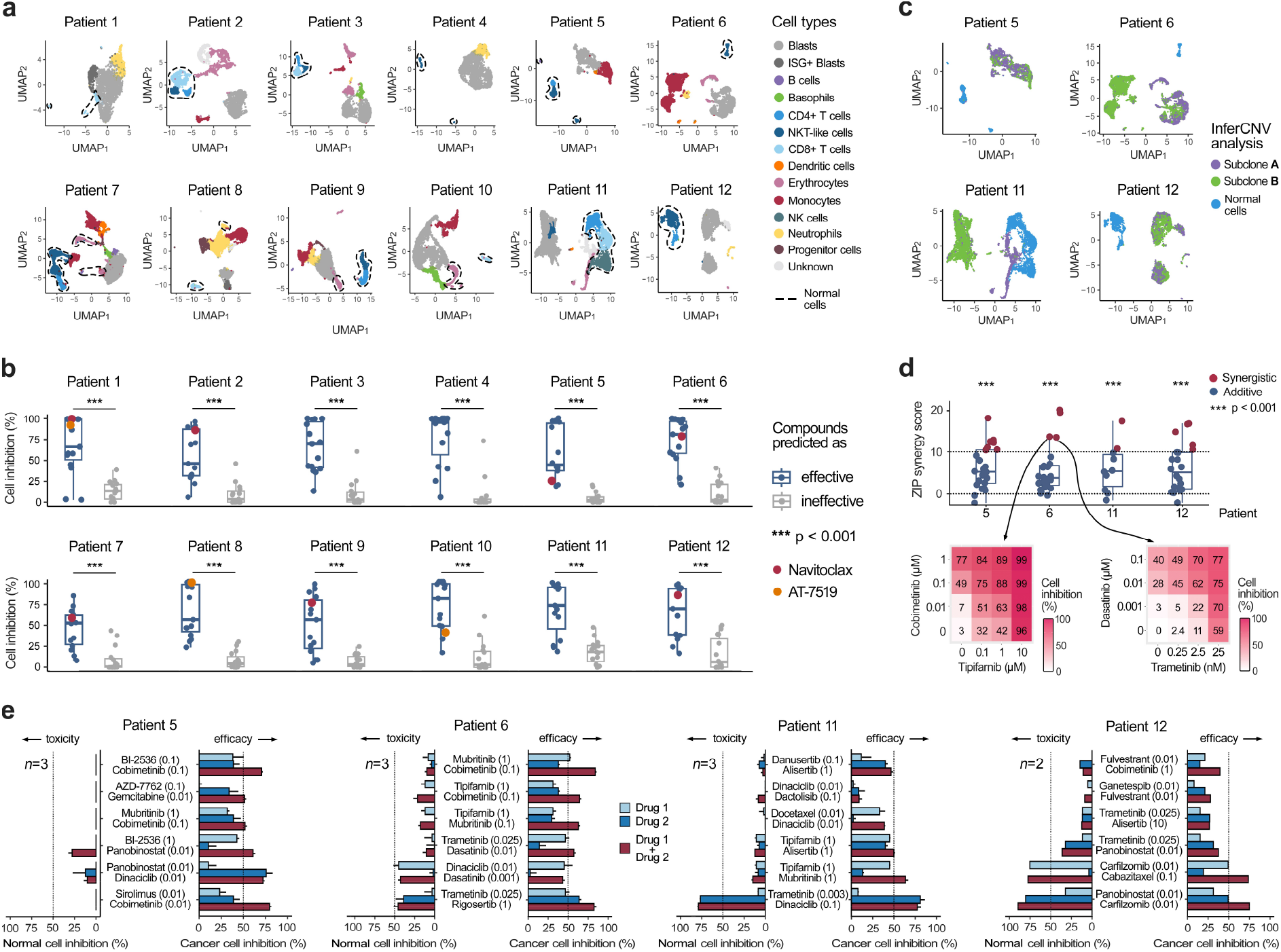
Experimental validation using bulk and cell population drug assays. (**a**) Identification of cell types using scRNA-seq profiles of complex bone marrow samples from 12 AML patients. (**b**) *Ex vivo* drug sensitivity differences between single-agent treatments predicted by ScTherapy to be either effective or ineffective in whole-well cell viability assays (p<0.001, Wilcoxon test). The colored points show two example drugs with highly variable responses across the patients. (**c**) Identification of genetically-distinct subclones from the 4 AML patient samples with enough cells for further experimental testing (**Suppl. Fig. 3** shows a detailed overview of genomic variation in these 4 samples). (**d**) All the model-predicted drug combinations exhibited either synergistic (ZIP>10) or additive effects (0<ZIP<10) in the whole-well combinatorial viability assay (p<0.001, Wilcoxon test; upper panel). Two examples of combinations with ZIP=13.6 and ZIP=13.5 as tested in multi-dose drug combination assays (lower panel). Interactive plots of the dose-response matrices for all the predicted combinations are provided at https://ianevskialeksandr.github.io/waterfall_plot.html. (**e**) Further validation of the top-combinations for the 4 patient samples using population-level flow cytometry assays in the same patient-derived cells. Toxic effects (left-hand bars) scored based on co-inhibition of normal cell populations, and therapeutic effects (right-hand bars) based on co-inhibition of malignant cells. The predicted effective doses are indicated in parentheses (μM), and the dotted vertical lines indicate 50% inhibition level. n, the number of replicate screens. Patient 12 has only two replicates due to limited cell availability and poor cell viability (35% live cells in DMSO wells after 72h).

To validate the model predictions, we first used data from single-agent cell viability assays, which confirmed that the model-predicted effective treatments led to significantly better cell inhibition efficacy *ex vivo*, when compared with the predicted ineffective treatments (P<0.0001, Wilcoxon test; **Fig. 2b**). Importantly, this improvement was not due to the model selecting higher drug concentrations for the effective-predicted treatments (**Suppl. Fig. 2a**). Most of the treatment predictions were uniquely identified for a single patient (**Suppl. File 1**), and the few shared treatments between patients, such as navitoclax and AT-7519, showed highly variable responses across the patient samples (**Fig. 2b**, the colored points).

Next, we predicted the most promising two-drug combinations for four AML patient samples with enough cells for further experimental testing. The patient-specific combinations were designed so that they would maximally co-inhibit two major leukemic subclones in each patient sample, while minimally co-inhibiting the patient-specific normal cells (**Fig. 2c**). Using a bulk combinatorial cell viability assay, we tested the predicted combinations in 4×4 dose-response matrices (https://ianevskialeksandr.github.io/scTherapyCombinations.html). Based on the zero interaction potency (ZIP) score, we confirmed that all the predicted combinations act either synergistically (ZIP>10), i.e., they jointly inhibit patient cells more than expected based on their individual effects (p<0.001, Wilcoxon test), or showed at least additive combination effects (ZIP>0; **Fig. 2d**). It has been argued that a combination efficacy is more important for clinically-effective combinations, while pharmacological synergy is not necessary for achieving improved clinical responses^13^.

After confirming the higher than expected combination effects in the bulk viability assays, we further tested a subset of top-6 patient-specific combinations for the four patient cases using high-throughput flow cytometry assays to quantify the differential inhibition between leukemic and normal cells in each patient sample *ex vivo*. Out of the 24 predicted drug combinations, 21 (88%) led to increased co-inhibition of the leukemic cells (**Fig. 2e**), when compared with the single-agent responses. For each patient case, we identified multiple combinations that led to higher than 50% co-inhibition of the blasts and other leukemic cells, suggested as potential treatment options. Importantly, only 3 of the 24 combinations (13%) showed >50% inhibition of T cells and other non-cancerous lymphoid cells, which should be discarded as potentially toxic combinations (i.e., trametinib-dinaciclib combination in Patient 11, and two carfilzomib combinations in Patient 12). Not only the effective treatments, but also the predicted doses of drugs in the combinations varied across the patients, indicating that an optimal balance between treatment efficacy and toxicity can be tailored for each patient.

### Application to ovarian cancer and validation in patient-derived tumor organoids

To investigate whether the prediction approach is applicable also to solid tumors, where large-scale *ex vivo* drug testing in primary patient cells is more challenging, we employed published scRNA-seq data^14,15^ from a cohort of HGSC patients (**Suppl. Table 2**)^10^. This patient cohort of metastatic tumors with poor responsiveness to standard chemotherapy represents a highly challenging case for personalized treatment identification. To distinguish cancer cells from non-cancerous cells, we used established tumor marker genes, including *PAX8, MUC16* (encoding CA-125) and *EPCAM*, collectively referred to as PAX8+ cells (**Online Methods**). To secure enough fibroblasts and other genetically normal cells for the treatment-selectivity assays, we integrated scRNA-seq data from three HGSC patients (**Fig. 3a**), based on the availability of cells from each patient for further experimental validation: PAX8+ tumor cells were available from Patient 1, whereas PAX8-normal cells were available from Patients 2 and 3. The expression of PAX8 tumor marker showed a clear separation between tumor cells and other cell populations in this “integrated patient” case (**Fig. 3b)**.

**Fig. 3.**
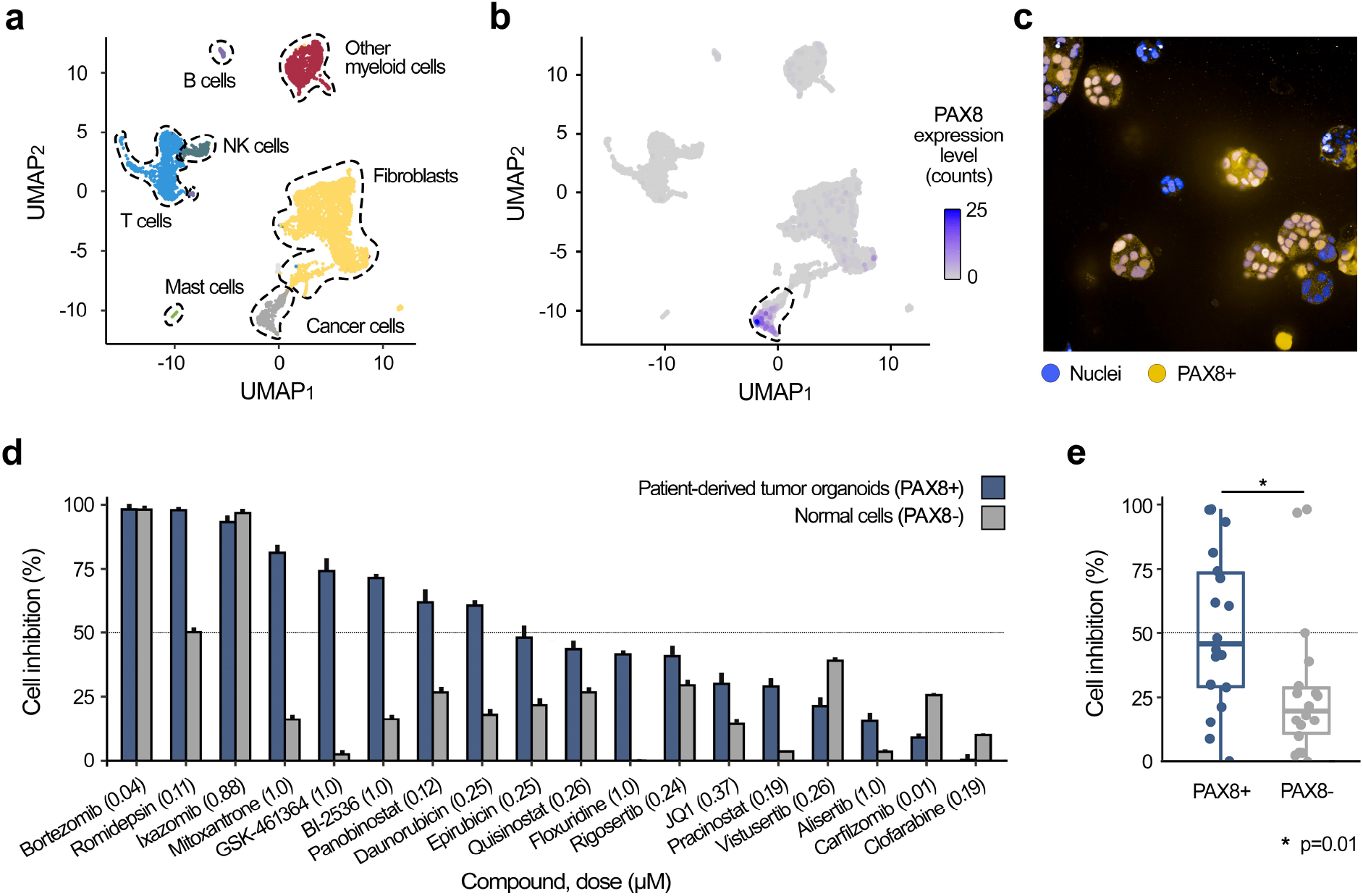
Experimental validation in ovarian cancer patient-derived tumor organoids. (**a**) UMAP projection of the combined scRNA-seq transcriptomic profiles from three HGSC patient samples, using standard Seurat integration workflow, where cell types were identified with ScType. UMAP plots for the individual patient samples are provided online https://ianevskialeksandr.github.io/figovfig.png. (**b**) Expression of the PAX8 marker, effectively separating tumor cells from the other cell populations. The expression of other PAX8+ markers is shown in **Suppl. Fig. 4**. (**c**) Representative immunofluorescent image of treatment-naive tumor organoids from Patient 1. (**d**) Cell inhibition differences between the patient-derived organoid cells and non-cancerous normal cells for the 18 predicted multi-targeting drugs. The predicted effective doses are indicated in parentheses (μM), and the dotted vertical lines indicate 50% inhibition level. The error bars represent SEM, based on three replicates of organoid treatments and curve-fitting approximation in PAX8-cells, respectively. (**e**) Statistical comparison of the treatment responses between PAX8+ and PAX8-cells (Wilcoxon test).

Due to a small proportion of patient-derived cancer cells, we predicted only multi-targeting monotherapies for this patient case, since combination treatments would require more cells for the organoid cultures. We tested the efficacy and selectivity of the predicted treatments on HGSC patient-derived tumor organoids^16^. The treatment-naive tumor organoids were developed exclusively from the cancer cells of a treatment-naive Patient 1 with omental metastasis (**Online Methods**), which displayed an elevated expression of PAX8 (**Fig. 3c**). Comparison of the treatment-induced viability changes in the organoid cells and fibroblasts showed that 8 of the 18 predicted treatments (44%) led to >50% inhibition of the PAX8+ tumor cells, and only 2 of the treatments (11%) showed >50% inhibition of the PAX8-cells (**Fig. 3d**; proteasome inhibitors bortezomib and ixazomib). In general, the predicted multi-targeting treatments led to significantly higher inhibition of the tumor cells than normal cells (p=0.01, Wilcoxon test; **Fig. 3e**). Interestingly, there was no correlation between the predicted treatment doses and PAX8+ or PAX8-cell inhibition effects (**Suppl. Fig. 2b**).

## Discussion

Advanced cancers are heterogeneous diseases, typically comprising at diagnosis more than 10^10^ cells, which very likely harbor therapy-resistant subpopulations^10,13^. This translates into a need for multi-targeting therapies for effective cancer cures that selectively co-inhibit the disease-driving cell populations. Our experimental-computational approach for personalized identification of multi-targeting treatments makes use of two recent advances: (i) the feasibility of scRNA-seq profiling in patient samples that does not only allow for the identification of malignant and non-cancerous cell populations, but can also elucidate the transcriptomic differences and hierarchies between these subpopulations; and (ii) the availability of large-scale transcriptomic and viability response profiles of cancer cell lines treated with thousands of single-agent perturbations. Taken together, this approach provides a clinically actionable and relatively fast means for predicting drug-dose combinations for individual patients, and compared to our earlier work^17^, it can be applied also for patients whose tumors are not easily amenable to drug testing (e.g. HGSC). The only input for the model is a count scRNA-seq data matrix of a given patient sample; the rest of the computational steps are either fully-automated or semi-automated (e.g., selection of the broad-level subclones based on visual analysis of the clonal evolutionary tree; see Step 4 in **Suppl. Fig. 1**). Nearly all of the predicted combinations exhibited positive synergy scores (96.3%), highlighting their potential for improved therapeutic efficacy and reduced toxicity by lowering the doses of single agents. Importantly, 85.7% of the predictions demonstrated low-toxicity to normal cells (<50% inhibition of non-cancerous cells); however, the flow cytometry and organoid drug response assays indicated that certain multi-target therapies (14.3%) excessively inhibited non-cancerous cells (e.g. proteasome and topoisomerase inhibitors), emphasizing the importance of experimental validation prior to clinical translation.

ScTherapy identifies individual drugs or their combinations that (i) induce normalizing transcriptomic changes in cell type-specific gene signatures of patient-specific DEGs (i.e., reverse clone-specific transcriptomic responses closer to the normal expression state), and (ii) exhibit selective cancer cell inhibition at the predicted effective dose (i.e., ensure differential inhibition between malignant and normal cells). The model outcome is a list of suggested treatments and concentrations for each patient sample, complemented with a confidence score that indicates the confidence of the LightGBM about the accuracy and reliability of each specific prediction. The quantitative performance evaluation (repeated cross-validation and experimental validations), together with the confidence scoring (conformal prediction), enables medical professionals to decide when and how to use the model to guide clinical decision making. By mapping the gene signatures to drug-target interactions networks, one can also explore potential biomarkers (e.g. patient-specific DEGs) that drive the selection of the best treatment regimens for individual patients (**Suppl. Fig. 5**). This provides additional insights into the rationale of the treatment recommendations for a given patient. ScTherapy model can also predict responses to custom compounds, hence facilitating the assessment of novel or less-studied compounds for their patient-specific efficacy. Furthermore, the model incorporates a user-defined drug-dose information, especially useful in cases where certain drugs or doses are clinically more relevant for a given cancer type. By dose restriction, one can further reduce the risk of toxic effects that often occur at higher doses, hence making the predictions clinically more relevant. Overall, our approach offers a systematic and flexible framework for predicting personalized drug-dose combinations that can be tailored to individual patient and tumor characteristics.

Traditionally, effective drug combinations have been identified either by empirical clinical testing^18^, or using high-throughput screening (HTS) in cell line panels *in vitro*, followed by target deconvolution and *in vivo* validation of the most relevant combinations and target mechanisms in animal models^19–21^. However, drug combination synergy is a rare and highly context-dependent event, affected by the inherent genetic and molecular variability between patients and within tumors, hence requiring combinations to be tested in large-scale screens and in various cellular contexts and genomic backgrounds^22^. This is beyond the scalability of *in vivo* models, and *in vitro* screening alone cannot identify combinations targeting specific cancer subclones, even if large enough cell line panels can to certain extent model the cellular heterogeneity and drug responses variability of tumors. This is important since a multi-targeting therapy that effectively inhibits cancer cells may also co-inhibit normal cells, rendering the treatment non-selective against malignant cells. In patient applications, it is therefore critical to identify cancer-selective combinations, rather than broadly active therapies that may lead to severe toxic effects. *Ex vivo* drug testing in primary patient cells, using either patient-derived 2D cell cultures or 3D organoids, strikes a balance between the *in vitro* and *in vivo* approaches^23,24^. However, even though flow cytometry and imaging-based *ex vivo* assays offer possibilities for drug response testing at a single-cell resolution, HTS of a larger number of drug combinations in multiple doses remains infeasible in scarce patient cells using these advanced assays^25–27^. Therefore, systematic methods to prioritize the most potential combinations to be tested in primary patient cells are needed.

Various machine learning (ML) methods have been developed to predict effective anticancer drug combinations, using multi-omics training data from large-scale screens in cancer cell lines and patient-derived samples. By surveying the existing ML methods^28^, we identified three critical areas of improvement for translational applications, where the aim is to prioritize among the massive number of potential drug-dose combinations those that show maximal therapeutic potential and minimal toxic effects for next phases of preclinical development. First, none of the existing methods were designed to predict selective drug combinations that target multiple cancer subclones in primary patient cells using merely single-cell transcriptomic data as input. This is important since multi-omics profiling and *ex vivo* drug testing in scarce primary patient cells is not yet practically feasible for many tumor types^24^. Second, most of the current methods either do not use any normal reference, and hence lack preclinical toxicity predictions, or use molecular or functional profiles from healthy individuals to de-prioritize toxic combinations, which may lead to non-selective combination predictions, due to high inter-individual molecular and phenotypic heterogeneity. Third, drug combination effects are not only patient-specific, but also highly dose-dependent, meaning that the same combination may show both synergistic and antagonistic effects at different dose windows^22^. To guide the eventual translation of the combination predictions, where lower dose combinations are often better tolerated by the patients, we argue that computational prediction methods need to provide dose-specific prediction of the responses. In this way, computational tools, such as ScTherapy, enable systematic *in-silico* screening of combination effects for translational applications to prioritize most potent combinations for further testing, among the massive number of potential drug-dose combinations^29^.

We predicted a higher number of targeted signal transduction inhibitors for the AML patients, compared to HGSC, which reflects underlying differences in the disease biology. AML cells often carry oncogenic mutations in signaling proteins, making the cells addicted to MAPK signaling^30^, which explains why MEK inhibitor combinations were identified for many of the AML patients. Similarly, PLK inhibition has been extensively studied in AML, and while PLK inhibitor combinations have shown promise in clinical development, they are also associated with complicated toxicities^31^. Therefore, even though the ScTherapy model identifies novel personalized multi-targeting treatments, the drug and target classes of the predicted combinations are well-studied in AML and HGSC. The novel concept is that the predictions are tailored to the molecular context of a given patient (or sample), which is expected to lead to better efficacy-safety balance at the level of an individual patient, rather than identifying broadly chemotoxic combinations that may lead to severe side effects in the non-matching subset of patients. Our functional precision medicine approach provides a streamlined, yet relatively precise approach to finding the right combinations of drugs and enhancing therapeutic potential through using both molecular and functional information.

### Limitations and future aims

Compounds from different drug and target classes can elicit varied phenotypic responses in the viability and transcriptional profiles. For instance, in contrast to other molecularly-targeted compounds, HDAC inhibitors often induce significant changes in the expression of multiple genes beyond their target proteins. Comparison of the expression and viability changes between cancerous and non-cancerous cells is expected to normalize out a part of such variability between drug classes. However, future studies are warranted to tailor input data not only to be patient-specific, but also drug class-specific by considering differences in binding affinities, phenotypic profiles and in treatment time points. Different disease models may also have differing growth dynamics. For instance, as opposed to the most conventional cell lines, organoid cells undergo less cell divisions during 7-day incubation. Therefore, some of the discrepancies seen between the model predictions (made using *in vitro* cell line data) and the experimental validations (made in *ex vivo* experiments) may stem from such variations between the 2D and 3D disease models and time points. Future studies are needed to identify the most predictive sources of *in vitro* and *ex vivo* data for patient-specific predictions in various cancer types. Additionally, with the emerging availability of large-scale morphological and proteomic response profiles in cancer cell lines and samples^32–35^, the ScTherapy approach could be extended to incorporate these and other phenotypic response measurements in the future. The approach is applicable also to selective targeting of other cell types or states, beyond the differential co-inhibition of cancer and non-cancerous cells.

## Materials and Methods

### Compiling a large-scale phenotypic response data for pre-training a LightGBM model

A comprehensive training dataset of large-scale phenotypic response profiles was created by merging data from three databases: Connectivity Map LINCS 2020^11^, PharmacoDB^12^, and PubChem^36^ (**Suppl. Fig. 1**, bottom part). These continuously expanding, publicly available databases allowed us to establish an extensive dataset that provides functional information on both viability and transcriptomic responses to increasing numbers of compounds. Details on the dataset used in the present study are outlined below. The Connectivity Map (CMap) LINCS 2020 is a reference database that houses gene expression response profiles of 12,328 genes measured in 240 cell lines across multiple doses and time points for 39,321 small-molecule compounds. Additionally, LINCS 2020 data includes paired control states for each perturbagen-cell line combination, enabling a comparison of the transcriptional changes before and after each treatment. To supplement our dataset, we leveraged information from PharmacoDB, a database that contains dose-response viability data for 56,149 drugs across 1758 cancer cell lines at multiple doses. For further analysis, we employed 10,303 overlapping compound-cell line pairs, which were common between 24 h transcriptional responses from CMap LINCS 2020 (passing quality control, i.e., qc_pass = 1) and PharmacoDB. For matching compounds between PharmacoDB and CMap LINCS 2020, we used compound identifiers, and for the cell line matching, we used cellosaurus IDs^37^. To extract structural information of the compounds, we used PubChem and RDKit (rcdk v3.6 and rcdklibs v2.3) to generate molecular fingerprints (ECFP4) from the SMILES representation of each common drug^38^.

The light gradient boosting machine (LightGBM) model was trained on a comprehensive dataset of 3,695 compounds tested at 1-35 doses in 167 cell lines. Drug-dose-cell line profiles (including transcriptomic response profiles, ECFP4 molecular fingerprints, and drug doses) were used as the model predictors, while the outcome variable is the inhibition percentage, derived from PharmacoDB dose-response viability data (**Suppl. Fig. 1**). The LightGBM model was trained using Bayesian Optimization, with a repeated cross-validation (three repetitions), and ten-fold inner cross-validation (CV). This ensures a robust and generalizable model for the patient applications. More specifically, the LightGBM model matches gene expression signatures (differentially expressed genes between the cancer and non-cancer cells) to the transcriptional responses to small molecules tested at different doses from LINCS 2020 to find the compounds that induce opposite transcriptomic changes. In the next step, the model identifies which compounds and doses most effectively inhibit cell growth, by extracting %inhibition responses for corresponding cell line-drug-dose triads from PharmacoDB. After examining tens of thousands of possible matches, the model provides a prediction of the most promising compounds and the effective dose. We also recommend including at least one dose-fold above and below the predicted dose in the experimental evaluation to delineate the most effective and least toxic drug dosage.

### Prediction of multi-targeting therapies using scRNA-seq data in AML patient samples

The experimental-computational prediction approach consists of the following five subsequent steps (**Suppl. Fig. 1**). These steps are described here for the AML case, and modifications to this pipeline in the HGSC case are described under section **Tailoring the experimental-computational approach to ovarian tumor patient samples**.

### Step 1: Longitudinal sampling

After obtaining informed consent, bone marrow aspirates were collected from patients diagnosed with acute myeloid leukemia (AML) at the Helsinki University Hospital (HUS). For this study, a total of 12 longitudinal samples (7 at diagnosis, 2 at relapse stage and 3 at refractory stage) were obtained and stored at the Finnish Hematology Registry and Clinical Biobank (FHRB). The protocols used for this study were reviewed and approved by the institutional review board in compliance with the Declaration of Helsinki^39^. The below steps 2-5 were repeated for each sample individually to provide a customized set of effective and low-toxic multi-targeting options for each patient individually by considering the intratumoral heterogeneity of cancer cells that is present not only at later stages of the disease or resistance development, but already at the diagnostic stage.

### Step 2: Single-cell data analysis

For the single-cell transcriptomic analysis, we processed the filtered gene-barcode matrix derived from 10X Genomics data using the ScType platform^40^, with Louvain clustering, as implemented in the Seurat version 4.3.0^41^. To filter out low-quality cells, we removed cells that had either a low or high number of detected genes and also cells that had more than 10% of mitochondrial UMI counts in the AML scRNA-seq data. The quality control (QC) criteria depend on the sample types; for instance, in HGSC organoids, 20% of mitochondrial UMI count cut-off was used^16^. Such QC cell filtering step is critical to exclude technical noise and thus to avoid biases in the downstream analysis. To normalize the gene expression levels, we utilized the LogNormalize method implemented in Seurat.

### Step 3: Identification of malignant and normal cells

Single-cell RNA sequencing profiles were used to identify malignant and normal cell clusters in each sample using three analytical tools, ScType^40^, CopyKAT^42^, and SCEVAN^43^. These tools were specifically selected for their ability to accurately classify and differentiate between malignant and normal cells in the given complex sample, eliminating the requirement for larger cohort samples.

### Step 3a: Cell type annotation

We utilized the ScType web-tool^40^ that enables fast, precise and fully-automated cell cluster annotation. ScType integrates cell type markers from the two most comprehensive resources for human cell populations, and classifies cells based on gene expression changes across clusters. We used ScType to assign a confidence score to each cell type annotation and each cluster, with high scores indicating a high level of confidence in the cell type annotation. Clusters with low scores were labeled as “Unknown” cell types based on the default ScType cutoff (score < number of cells in the cluster divided by 4). In addition, we visually analyzed previously established marker genes for blasts, including CD33, CD34, CD38, PROM1, ENG, CD99 and KIT (17)^17^, on the UMAP space and calculated the proportion of the blast cells in each patient sample to gain a better understanding of the distribution of leukemic cells. This resulted in a Seurat object that includes cell clusters and their corresponding annotations.

### Step 3b: Detection of aneuploid cells

To further classify cell populations as normal or malignant, we developed an ensemble approach that utilizes multiple methods to generate a confident classification. The first method is a marker-based approach, which involves carefully filtered cell markers from CellMarker2.0 database^44^, and then using these as a custom marker dataset for ScType to identify normal and malignant cells. The second approach uses CopyKAT^42^, a Bayesian segmentation-based method, with default parameters and known normal cells (T cells in the AML case^45^) as a baseline to estimate copy number alterations (CNA). The third method is SCEVAN, with the non-cancerous control cells used as input, which employs a Mumford and Shah energy model to distinguish normal and malignant cell states^43^. The use of CNA estimation based approaches allows us to classify malignant cells while taking into account overall variability within normal cells. We then constructed a majority vote based on the combined results of these tools to confidently identify both normal and malignant cell clusters. To further validate our approach, we superimposed the ensemble predictions onto the UMAP space and compared them with the cell-type information obtained from ScType. By integrating cell type and normal/malignant annotations from ScType, with ploidy information from CopyKAT and SCEVAN, we identified clusters of cells as either normal or malignant. Our ensemble approach accounts for variability within normal cells and therefore minimizes the risk of misclassification.

### Step 4: Identification of genetically distinct subclones and visualizing clonal lineages

After successfully identifying normal and malignant cell clusters, we used inferCNV^46^ to infer large-scale copy number variations, such as gains or deletions of whole chromosomes or segments from the scRNA-seq data. The input for the inferCNV analysis included the known non-cancerous cells identified in Step 3, genomic locations, cell type annotations, and the scRNA-seq count matrix data. CNVs were inferred using the Hidden Markov Model (HMM) approach implemented in the 6-state i6 HMM model (https://github.com/broadinstitute/infercnvApp/blob/master/inst/shiny/www/Infercnv-i6-HMM-type.md). In accordance with the inferCVN guidelines in the document “Using 10X data’’ section (https://github.com/broadinstitute/infercnv/wiki/infercnv-10x), we adjusted the “cutoff” parameter from 1 to 0.1, and subsequently computed the CNV profiles from the scRNA-seq expression counts. To explore the subclonal structures, we used the “subcluster” method on the HMM predicted CNVs.

After identifying the genetically distinct subclones, we used Uphyloplot2^47^ to visualize intra-tumoral heterogeneity and clonal evolution using the CNV calls from the inferCNV 6-state HMM “subcluster” method and its “.cellgroupings” file. We note that the resultant evolutionary tree does not follow a molecular clock; rather, the branch length is proportional to the percentage of cells in the subclone, hence providing information about which subclone dominates the tumor mass. Next, two broad-level subclones detected from the evolutionary tree were identified and, along with normal cells, overlaid on a UMAP projection for further analysis. To quantify gene expression differences between the normal cells (identified in Step 3) and the broad-level subclones (identified in Step 4), log-fold change values and determined significance levels via the nonparametric Wilcoxon rank-sum test, applied in Seurat 4.3.0 using the FindMarkers command.

### Step 5: Predictive modeling of multi-targeting therapies

When applied to patient samples, the subclone-specific differentially expressed genes (DEGs) were used as input for the pre-trained LightGBM model to predict single-agent cell inhibition percentages for each compound-dose pair in the particular patient cells. This allows us to take into account both the intratumoral and intertumoral heterogeneity, as captured by the scRNA-seq profiles of the patient samples. Our prediction approach is highly flexible and can be used in two ways: first, by utilizing a predefined set of drug-dose pairs for predictions, or second, by customizing the analysis with additional input of new drug structures (ECFP4 fingerprints) and/or specific doses of interest.

As any ML model predictions inherently come with some degree of uncertainty, we used conformal prediction (CP) to eliminate low-confidence predictions and improve the prediction accuracy^48^. CP generates confidence intervals for each prediction by measuring uncertainty based on repeated CV residuals. Predictions with a nonconformity score <0.8 were excluded, thereby ensuring inclusion of only confident and accurate predictions. In addition, to ensure that our model returns clinically more relevant predictions, we imposed a 1 μM dose maximum when utilizing the pre-defined set of drug-dose pairs. High drug doses, even though potentially increasing cancer cell inhibition, may also inhibit normal cells, hence compromising the selectivity of targeted agents^49^. By using such a dose restriction, we ensured the selectivity of targeted agents returned by the model and minimized the risk of toxic effects, making our predictions more clinically actionable. We applied this approach to each subclone, hence generating a set of drug-dose-response tuples for the experimental validation.

### Retrospective testing of the model predictions in single-agent data from AML patients

To validate the performance of our model, we first used existing data from bulk drug response assays, available for the 12 patient samples from previous studies^39^. For the single-agent response testing, 20 μl of fresh AML cell (approximately 10,000) suspension in mononuclear cell medium was added per well to pre-drugged plates with 10-fold dilution series of five concentrations, and the whole-well cell viability was measured with CellTiter-Glo (CTG; Promega) in duplicate, as previously described^30,39^. After 72 h of incubation at 37°C and 5% CO2, cell viability of each well was measured using the CTG luminescent assay and a PHERAstar FS (BMG Labtech) plate reader. The percentage inhibition was calculated by normalizing the cell viability to negative control wells containing 0.1% dimethyl sulfoxide (DMSO), and positive control wells containing 100 μM cell killing benzethonium chloride (BzCl). Notably, these existing single-agent response data were not used in the model training, and were only employed retrospectively to test the accuracy of the model to predict effective monotherapies. Since the whole-well assay is not a cell population-specific assay, we performed this validation using the differentially expressed genes (DEGs) between the malignant cell types and normal cells to generate single-agent predictions for each patient sample. Subsequently, we matched the drugs and doses predicted by the model to the available patient-specific cell viability dose-response data (**Fig. 2b**).

### Prospective testing using whole-well and flow cytometry assays in the AML patient cells

The patient-specific predicted combinations were first tested on the bone marrow mononuclear cells of each patient in a 4 × 4 dose-response matrix using the bulk CTG viability assay, similarly as before^17^. The combination synergy in the experimental validations was quantified using ZIP model^50^, calculated based on the dose region around the predicted effective dose of each compound in the combination.

Cell population-specific drug combination effects in primary AML patient samples were assessed by high-throughput flow cytometry assay, as described previously^17,51^. Briefly, the compounds were dissolved in 100% dimethyl sulfoxide and dispensed on conical bottom 384-well plates (Greiner) either as single agents or combinations using an Echo 650 liquid handler (Beckman Coulter). Cryopreserved bone marrow mononuclear cells were thawed and suspended in 12.5% HS-5 derived conditioned medium, and 2-3x10^4^ live cells were seeded with a MultiFlo FX.RAD (BioTek) to 384 well-plates, followed by incubation for 72 h at 37°C and 5% CO^2^. To profile the cell subpopulation responses, the cells were stained with BV785 Mouse Anti-Human CD14 (Biolegend), VB515 Recombinant Anti-Human CD56 (Miltenyi), and following antibodies from BD Biosciences; V500 Mouse Anti-Human CD45, BV650 Mouse Anti-Human CD19, PE-Cy7 Mouse Anti-Human CD3, PE Mouse Anti-Human CD34, BV421 Mouse Anti-Human CD38 and APC Mouse Anti-Human CD117, together with APC-Fire 750 Annexin V (Biolegend) and DRAQ7 (BD Biosciences). The cells were analyzed with an iQue3 flow cytometer (Sartorius). Remaining live cells after drug treatments were gated using Forecyt (Sartorius). Briefly, cell singlets were identified on the basis of FSC-A (forward-scattered area) versus FSC-H ratio, and live cells were identified by excluding annexin V-and DRAQ7-positive cells, followed by identification of leukocytes (CD45+). Further characterization was done for NK cells (CD56+CD3-), leukemic blasts (CD34+ and/or CD117+) leukemic stem cells (CD34+CD38-), monocytes (CD14) and T/B-cells (SSC-A and CD3/19) from the leukocytes.

### Tailoring the experimental-computational approach to ovarian tumor patient samples

To differentiate between cancer and non-cancerous cells in ovarian cancer patient scRNA-seq data, we utilized a panel of established marker genes, including PAX8, CA125, MUC16, WFDC2, and EPCAM, collectively referred to as PAX8+ cells; PAX8 is expressed in 80–96% of high-grade serous ovarian cancer (HGSC) tumors (**Suppl. Fig. 4**)^52,53^. Our initial analysis focused on the HGSC Patient 1 sample, selected due to the availability of both scRNAseq data and viable cells for experimental validation. Due to the small proportion of PAX8+ malignant cells detected in the scRNAseq data, we opted to predict only single-agent therapies as opposed to combination therapies. However, during the validation phase, the PAX8-fibroblasts of Patient 1, serving as normal controls, died. This led us to integrate this sample with two other HGSC Patient 2 and 3 samples, which had readily available PAX8-cells. The integration was achieved using the standard Seurat workflow, and the cell types were assigned using ScType. Both combined PAX8+ and PAX8-cell populations were visualized using Seurat “FeaturePlots”. We used an average of previously-measured responses of PAX8-cells from patient 2 and 3 samples (serving as combined ovarian-sample normal controls) to 372 compounds overlapping with the LINCS 2020 compounds.

### Prospective testing in ovarian tumor organoids and drug response assays

To predict the compounds that specifically target and eliminate cancer PAX8+ cells, while sparing PAX8-cells, we utilized the differentially expressed genes (DEGs) from the comparison between PAX8+ and PAX8-cells in the scRNA-seq data. These DEGs were used as input for the pre-trained LightGBM model. Among the predicted 372 compound responses (that overlapped with drugs tested on PAX8-cells), we selected the top-20 most effective compounds, and removed two with low confidence, hence resulting in 18 predicted agents. Subsequently, we validated the efficacy of these compounds in PAX8+ tumor organoids and compared the results, as shown in **Fig. 3b** (3 replicates).

Ovarian cancer organoids were established and characterized as previously^16^, and propagated in BME-2 matrix droplets in the sample-specific growth medium. The organoid cultures consisted only of cancer cells as judged by whole-genome sequencing and copy number variation analysis. For the organoid drug sensitivity testing, the organoid cultures were trypsinized to obtain the single-cell suspension. The cells were resuspended in the fresh gel, dispensed to 384-well Ultra-Low Attachment microplates (#4588, Corning) at 10^3^ cells per well in 10 µl of the matrix, and covered with 60 µl of growth medium containing 5 µM ROCK inhibitor to facilitate the organoids formation. After 6 days, the medium was exchanged to 30 μl/well of the ROCK inhibitor-free growth medium. Drug testing was performed as described above for single-agent AML sample testing, with the following modifications. The tested compounds (10-fold dilution series of five concentrations), vehicle (DMSO), or positive control compounds (100 μM benzethonium chloride or 10 μM staurosporine) were transferred to the wells using Echo 550 acoustic dispenser (Labcyte). The organoids were incubated with drugs for 7 days in the humidified incubator at 37°C and the viability was assessed using CellTiter-Glo 3D Cell Viability Assay (#G9683, Promega) using a SpectraMax Paradigm microplate reader (Molecular Devices) after 5 min of agitation and 25 min of incubation at room temperature, as indicated by the manufacturer. The PAX8-negative cells from the ovarian tumor samples were expanded in RPMI-1640 medium, supplemented with 2 mM glutamine, 1% Pen/Strep and 10% fetal bovine serum (Gibco). Drug testing was performed as above. The culture was trypsinized, resuspended in fresh medium and seeded at 1000 cells in 25 μl of medium per well in pre-drugged 384-well microplates (#3864, Corning). After 7 days of the drug treatment, the viability was measured using the CellTiter-Glo 2.0 (Promega).

### Ethical approvals

AML patient samples and data were collected with signed informed consent in accordance with the Declaration of Helsinki (Ethical Committee Statement 303/13/03/01/201, latest amendment 7 dated June 15, 2016. Latest HUS study permit HUS/395/2018 dated February 13, 2018). HGSC patients participating in the study gave their informed consent, and the study was approved by the Ethics Committee of the Hospital District of Southwest Finland (ETMK 145/1801/2015).

## Data and code availability

- The raw single-cell RNA sequencing data for the AML Patients 5, 6 and 12 have been made available at the European Genome-phenome Archive (EGA) under the code (pending).
- The previously published single-cell RNA-seq data of the remaining 9 AML patient samples are available at EGA. The raw single-cell RNA sequencing data for the AML Patients 2, 3, 8 and 10 are available under the code EGAS00001004614^17^, and for the AML Patients 1, 4, 7, 9 and 11 under the code EGAS00001004444^54^.
- The previously published single-cell RNA-seq data of the 3 HGSC patient samples are available at EGA. The raw single-cell RNA sequencing data for the HGSC Patient 1 and Patient 2 are available under the code EGAS00001005010^14^, and raw data for the HGSC Patient 3 under the code EGAS00001005066^15^.
- The R codes for reproducing and making new patient-specific predictions are available at GitHub: https://github.com/kris-nader/scTherapy

## Acknowledgements

We are most grateful to the patients and their families for participating in the studies. The CTG and flow cytometry drug assays were carried out at the FIMM High Throughput Biomedicine Unit, which is hosted by the University of Helsinki and supported by HiLIFE and Biocenter Finland. Single-cell RNA sequencing of AML patient samples was carried out by the FIMM Single Cell Analytics Unit. The authors thank Minna Suvela for preparation of the AML patient samples for scRNA-seq, Dr. Juho Miettinen for help with the AML single-cell data, Dr. Imre Västrik for help with the AML patient data, Dr. Esa Pitkänen for help with the genomic analyses, Dr. Laura Gall-Mas for help with the HGSC organoid cultures, as well as Matias M. Falco and Yingjia Chen for their help with the ovarian cancer single-cell data.

