## Supplementary information for "Single-cell transcriptomes identify patient-tailored therapies for selective co-inhibition of cancer clones"

[1]Institute for Molecular Medicine Finland (FIMM), HiLIFE, University of Helsinki, Helsinki, Finland
[2]Biotech Research and Innovation Centre (BRIC), University of Copenhagen, Copenhagen, Denmark
[3]Foundation for the Finnish Cancer Institute (FCI), Helsinki, Finland
[4]Research Program in Systems Oncology, Research Programs Unit, Faculty of Medicine, University of Helsinki, Helsinki, Finland
[5]Department of Hematology, Helsinki University Hospital Comprehensive Cancer Center, Helsinki, Finland
[6]iCAN Digital Precision Cancer Medicine Flagship, University of Helsinki and Helsinki University Hospital, Finland
[7]Institute for Cancer Research, Department of Cancer Genetics, Oslo University Hospital, Oslo, Norway
[8]Oslo Centre for Biostatistics and Epidemiology (OCBE), Faculty of Medicine, University of Oslo, Oslo, Norway
[#]These authors contributed equally to this work
*Correspondence: Tero Aittokallio, PhD, Institute for Molecular Medicine Finland, FIMM, P.O. Box 20 (Tukholmankatu 8), FI-00014 University of Helsinki, Helsinki, Finland. Tel: +358 50 3182426,

### Supplementary Information

- **Supplementary Tables 1-2**
- **Supplementary Figures 1-5**
- **Supplementary File 1.** AML patient-specific drug response predictions and experimental validations across 12 patient samples

**Supplementary Table 1 | AML patient samples used in the study.**

| Pt | Disease stage | Age | Sex | scRNA-based blast (%) | Clinical morphological blast (% | Diagnosis (ICD-O) | FAB type | ELN22 risk class | Potential driver mutations | Treatment history | No. cells after QC[a] |
|---|---|---|---|---|---|---|---|---|---|---|---|
| 1 | Diagnosis | 68 | F | 79.3 | 80 | AML C92 | NA | Favorable | NPM1,DNMT3A,NRAS,IDH1,EML4 | - | 2483 |
| 2 | Diagnosis | 70 | F | 38.1 | 70 | AML C92 | M1 | Favorable | NPM1, TET2,USP8 | Azacitidine | 2874 |
| 3 | Refractory | 71 | F | 56.7 | 42 | AML C92 | M1 | Favorable | NPM1, TET2, HDAC 1,2,7 | Azacitidine-Venetoclax | 2421 |
| 4 | Diagnosis | 21 | F | 80.3 | 91 | AML C92 | M1 | Favorable | RANBP2,NPM1,IDH1,FLT3 | - | 2140 |
| **5** | **Relapse** | **71** | **F** | **54.4** | **26** | **AML C92** | **M5** | **Intermediate** | **LZTR1,AR,ICOSLG** | **Hydroxyurea, Cytarabine** | **2365** |
| **6** | **Relapse** | **46** | **F** | **40** | **14** | **AML C92** | **M1** | **Intermediate** | **FLT3,PTPN11,TP53BP1** | **Cytarabine, Cytarab-Amsacrine, Clofarabine-Cytarabine-Etoposide, Azacitidine,Azacitidine-Lenalidomide** | **5677** |
| 7 | Diagnosis | 75 | M | 40.8 | 36 | AML C42 | NA | Adverse | RUNX1,BCORL1,PTPN11 | - | 9340 |
| 8 | Refractory | 68 | M | 22.8 | 40 | AML C92 | NA | Adverse | DNMT3A, ERG, U2AF1, BCOR | Cytarabine-Idarubicin, Azacitidine | 4461 |
| 9 | Diagnosis | 68 | F | 50 | 70 | AML C92 | M1 | Adverse | BCOR, IDH2 | - | 3921 |
| 10 | Diagnosis | 35 | M | 65.5 | 65 | AML C92 | M2 | Adverse | WT1, CCND2, CEBPA | Cytarabine-Idarubicin, Lenalidomide | 3111 |
| **11** | **Diagnosis** | **74** | **M** | **42.1** | **32** | **AML C92** | **M2** | **Intermediate** | **VAV1** | **-** | **5697** |
| **12** | **Refractory** | **71** | **M** | **65.3** | **65** | **AML C92** | **M2** | **Adverse** | **MN1,MAP2K2,ETV6,FOXP1** | **Aspirin, Anagrelide** | **3610** |

[a]Quality control: filtering out low-quality cells (see Step 2 in the Methods section).
**Boldfacing** indicates patient samples with enough cells for experimental validation.

**Supplementary Table 2 | HGSC patient samples used in the study.**

| Patient ID | Treatment | Stage[a] | Anatomical location of scRNA-seq | Cell types | No. cells before QC | No. cells after QC |
|---|---|---|---|---|---|---|
| Patient 1 | Treatment naïve | IIIC | Omentum | PAX8+ | 1921 | 1743 |
| Patient 2 | Treatment naïve | IIIC | Peritoneum | PAX8- | 711 | 564 |
| Patient 3 | Treatment naïve | IIIC | Ascites | PAX8- | 3666 | 2399 |

[a]Stage IIIC (FIGO staging): The cancer is in one or both ovaries or fallopian tubes, or there is primary peritoneal cancer, and it has spread or grown into organs outside the pelvis. The deposits of cancer are larger than 2 cm (about 3/4 inch) across and may be on the outside (the capsule) of the liver or spleen (T3c).

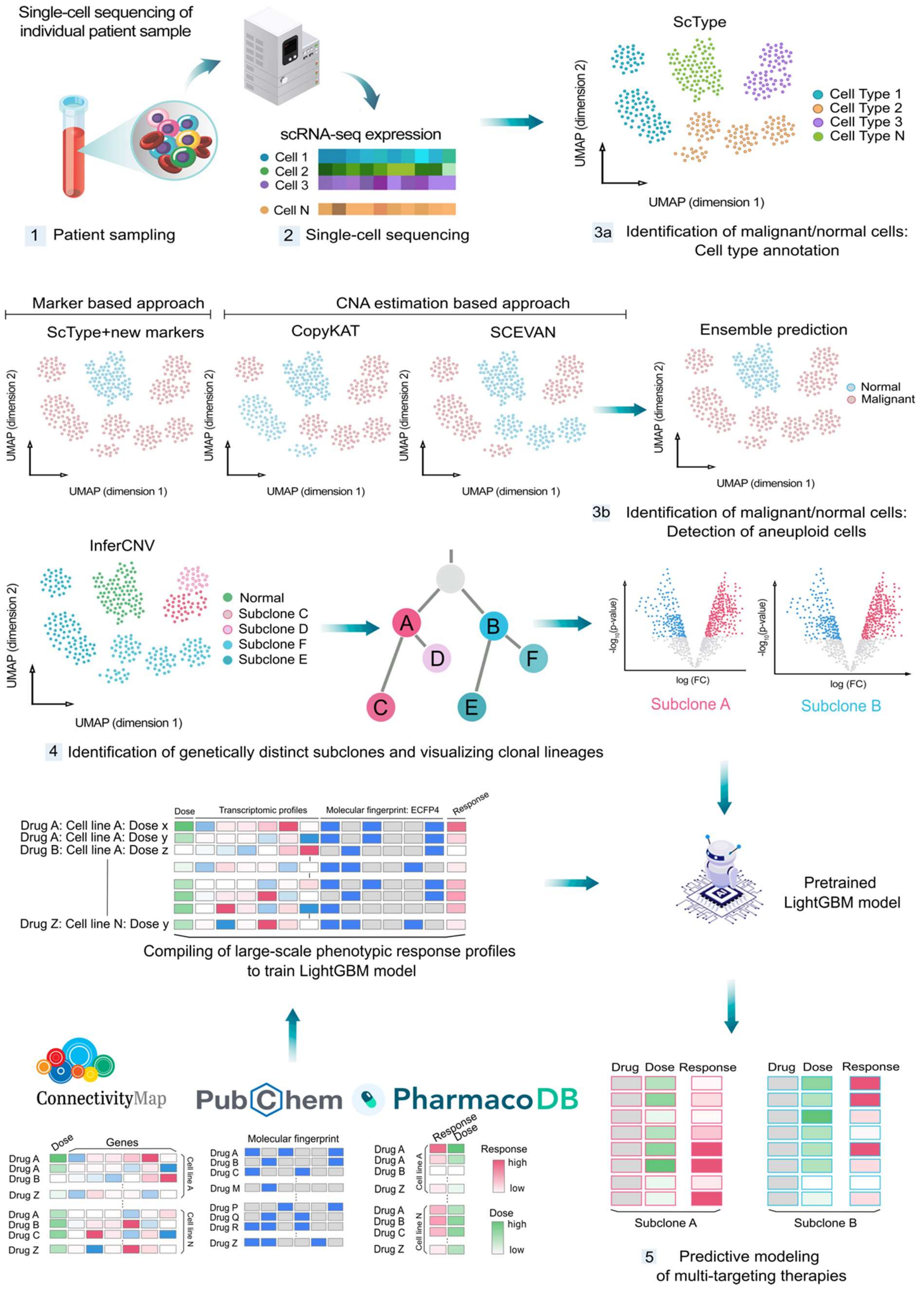


**Supplementary Fig. 1 | Schematic workflow of the experimental-computational approach to predicting multi-targeting treatments for an individual AML patient/sample.** See Methods for details of steps 1-5.

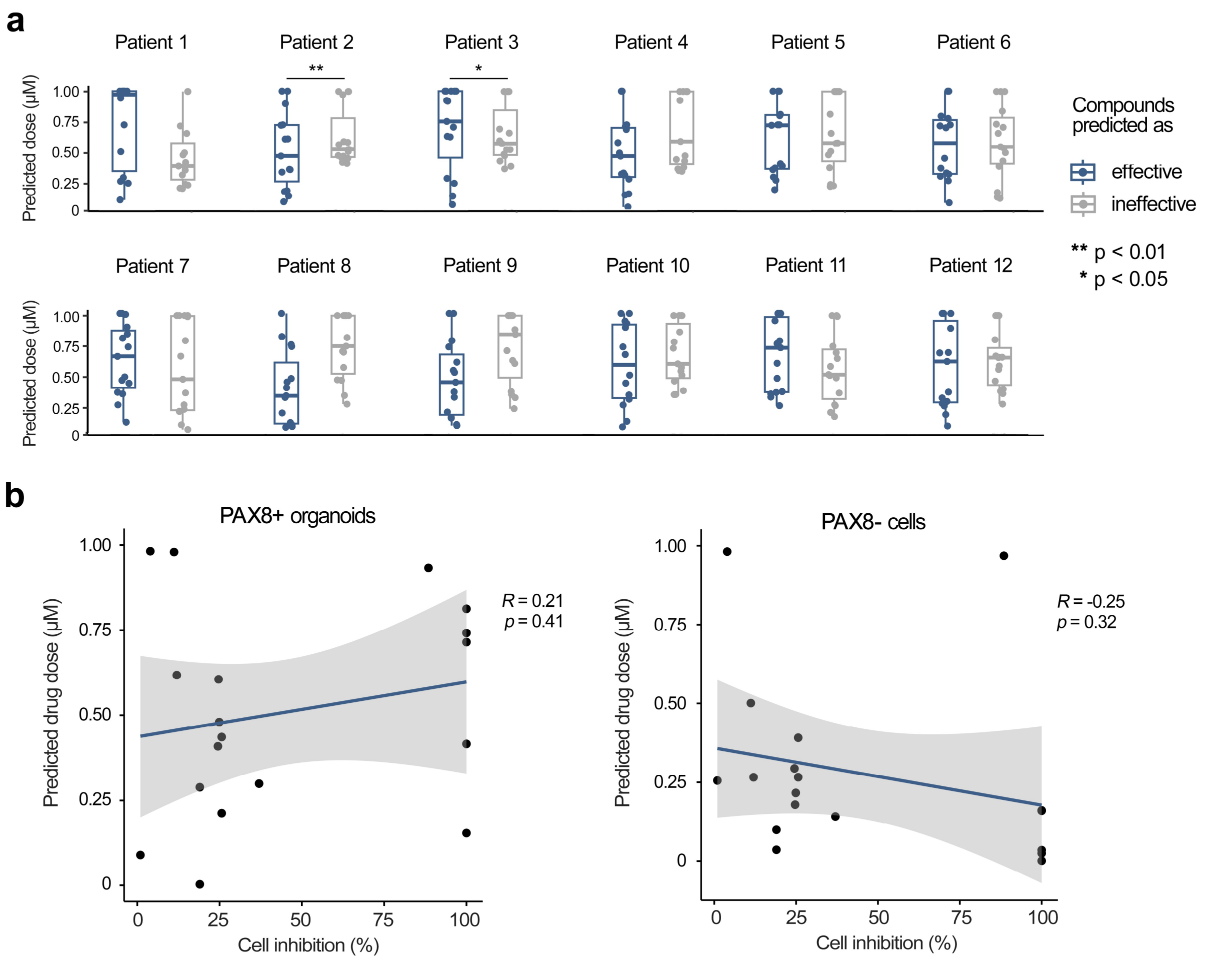


**Supplementary Fig. 2 | Drug doses of the predicted single-agent treatments.** (**a**) Drug doses (µM) for single-agent treatments predicted by the model to be either effective or ineffective for 12 AML patients (Wilcoxon test). In general, there is no significant difference in the doses between drugs predicted to be effective or ineffective. **(b)** Correlation between the predicted drug doses and the measured PAX8+ or PAX8- cell inhibition across the 18 treatments for the integrated HGSC case. There is no significant correlation based on correlation test.

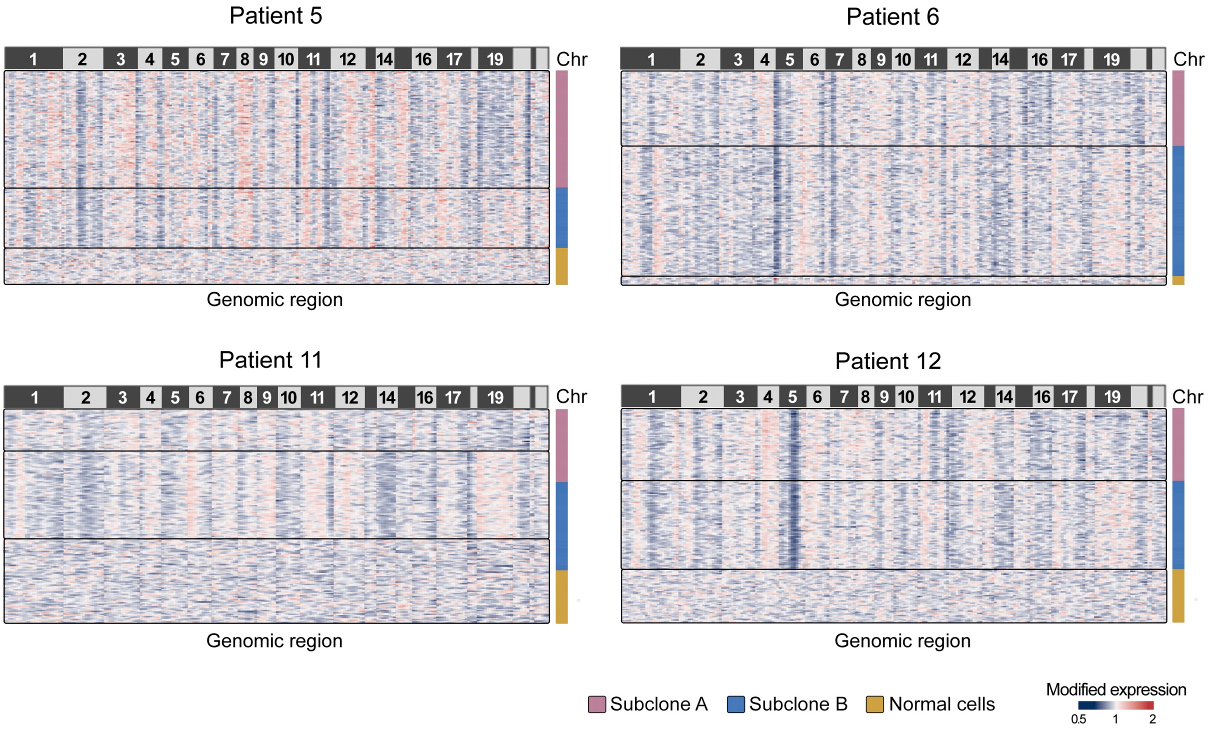


**Supplementary Fig. 3 | InferCNV copy number variation analysis of malignant cells from the four AML samples.** The heatmaps show a graphical representation of the CNV across the genomic regions, denoted as chromosomes (Chr) at the top of each heatmap. Shades of blue indicate lower levels of modified gene expression, suggesting genomic loss, while shades of red indicate higher levels of modified gene expression, indicative of genomic gain.

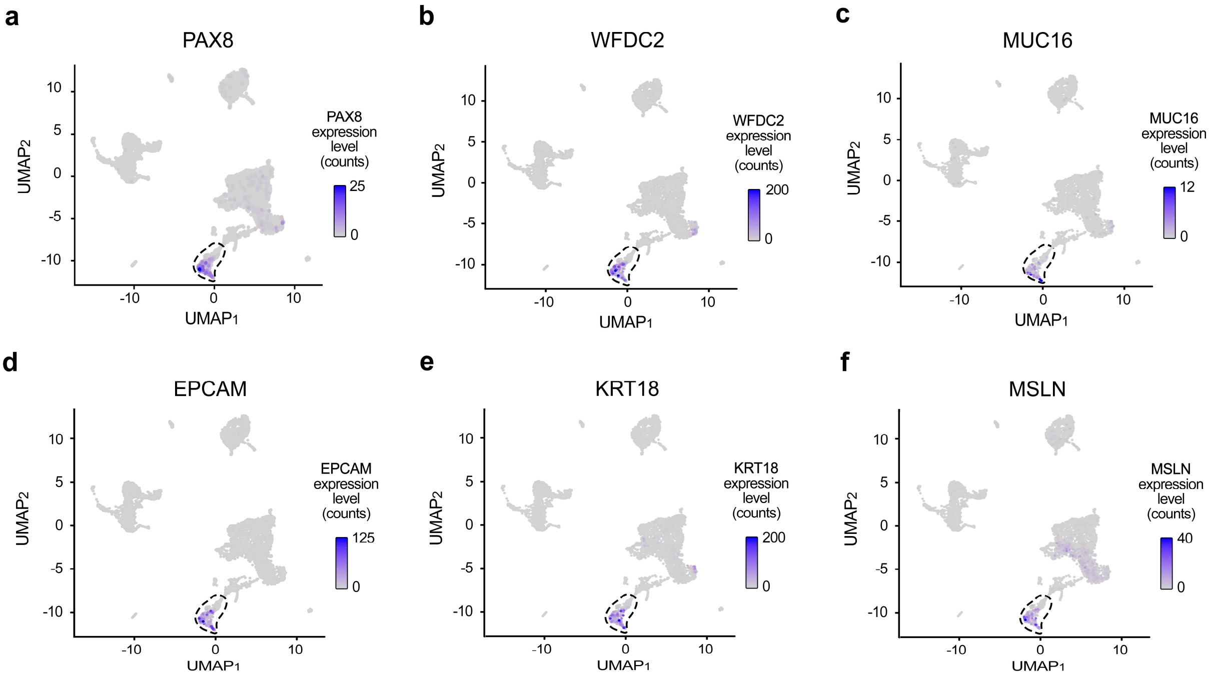


**Supplementary Fig. 4 | Expression of the PAX8+ marker genes for the detection of ovarian cancer cell populations.** (**a**) PAX8, (**b**) WFDC2, (**c**) MUC16, (**d**) EPCAM, (**e**) KRT18, and (**f**) MSLN. These markers were utilized for the detection of cancer cell populations in the HGSC Patient 1 (highlighted with dashed circles).

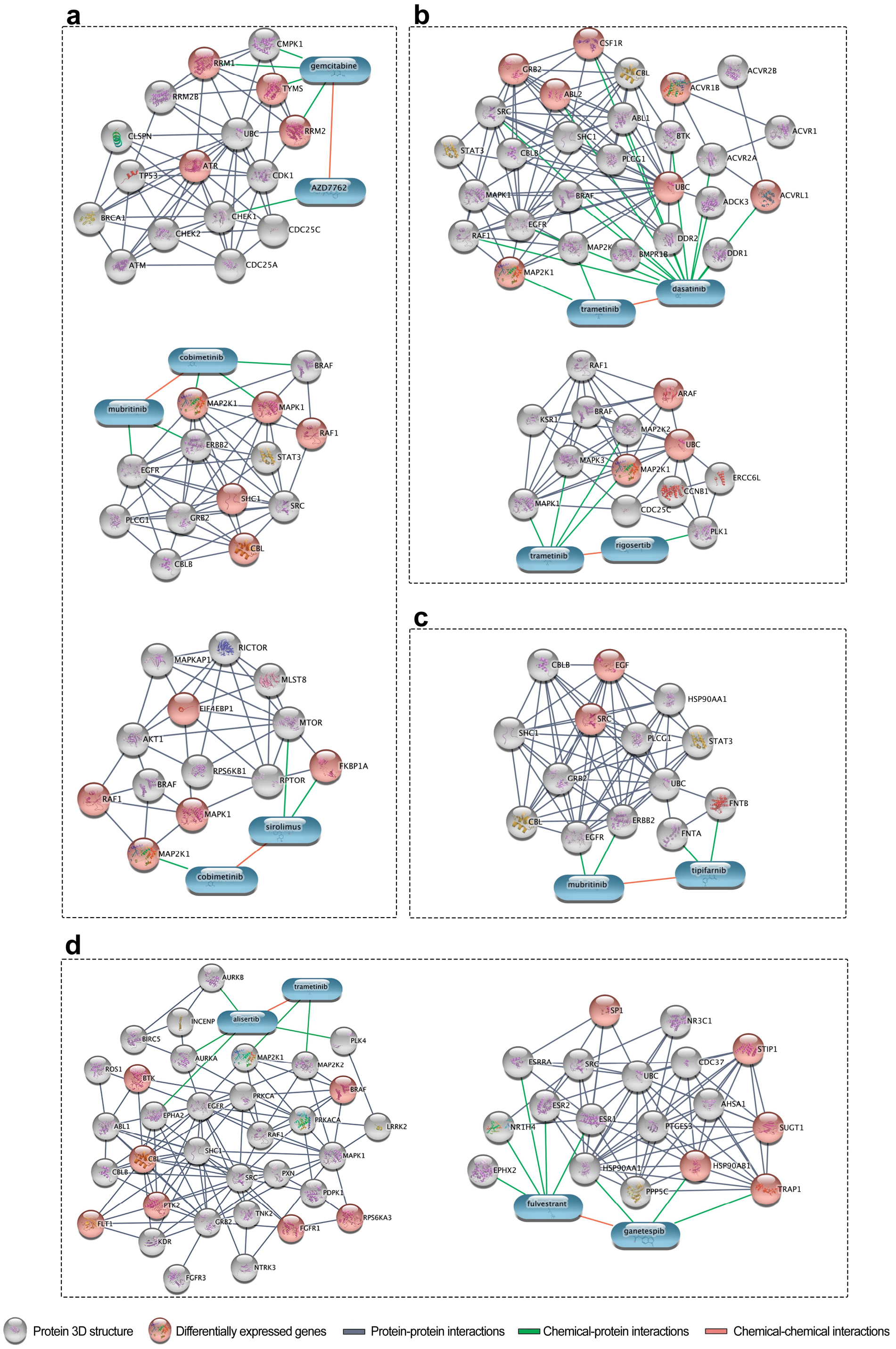


**Supplementary Fig. 5 | Interaction networks for the predicted and experimentally validated AML patient-specific combinations.** Representative drug combinations, where multiple drug targets were identified as differentially expressed genes between normal and malignant cells for the 4 AML patients: (**a**) Patient 5, (**b**) Patient 6, (**c**) Patient 11, and (**d**) Patient 12. The protein nodes in the networks include the nominal and potent off-targets of the compounds in the combinations, along with differentially expressed genes in the target pathways that may partly explain the observed combination effects in the particular patient cases. The network visualizations were done using the STITCH web-tool (Szklarczyk, D. et al. STITCH 5: augmenting protein-chemical interaction networks with tissue and affinity data. Nucleic Acids Res. 44, D380-384 (2016).
